# Tychus: a whole genome sequencing pipeline for assembly, annotation and phylogenetics of bacterial genomes

**DOI:** 10.1101/283101

**Authors:** Christopher Dean, Noelle Noyes, Steven Lakin, Pablo Rovira-Sanz, Xiang Yang, Keith Belk, Paul S. Morley, Rick Meinersmann, Zaid Abdo

**Author notes:** The authors wish it to be known that in their opinion, the first three authors should be regarded as joint first authors.

## Abstract

**Summary:** Tychus is a tool that allows researchers to perform massively parallel whole genome sequence (WGS) analysis with the goal of producing a high confidence and comprehensive description of the bacterial genome. Key features of the Tychus pipeline include the assembly, annotation, alignment, variant discovery and phylogenetic inference of large numbers of WGS isolates in parallel using open-source bioinformatics tools and virtualization technology. All prerequisite tools and dependencies come packaged together in a single suite that can be easily downloaded and installed on Linux and Mac operating systems.

**Availability:** Tychus is freely available as an open-source package under the MIT license, and can be downloaded via GitHub (https://github.com/Abdo-Lab/Tychus).

## 1 Introduction

The zeitgeist of the genomics era has been defined by the accessibility and affordability of next-generation sequencing platforms, which are capable of producing large amounts of data for genomics research. Whole genome sequencing (WGS) allows for the interrogation and analysis of complete genomes and can be applied to a wide range of organisms including plants, animals, bacteria, and viruses. Recently, WGS methods have been used in the domain of public health in order to better understand and track foodborne illnesses caused by bacterial pathogens. These efforts have led to the development of open-source reference databases and foodborne outbreak detection and investigation initiatives (Barkley et al., 2016; see also Hu et al., 2016). However, the accuracy of WGS for bacterial pathogen identification and outbreak analysis relies not only on the methods used to isolate, extract, and sequence the bacterial DNA, but also on the bioinformatics and statistical analyses applied to the resulting sequence data. In addition, as WGS datasets expand exponentially, there is a critical need for widely-accessible and computationally-efficient analytic pipelines. To this end, we present Tychus, an open-source, easy-install, user-friendly and rapid software tool for performing large-scale, parallel bacterial sequence analysis that produces a high confidence and comprehensive description of the bacterial genome.

## 2 Methods and Implementation

Tychus is written using Nextflow (Tommaso *et al*., 2015), a parallel DSL workflow framework and integrated with Docker (Merkel, 2014), a software containerization platform that resolves the installation and configuration issues of the many open-source bioinformatics tools utilized throughout the pipeline. Tychus is split into two complementary modules (see Figure 1): assembly and alignment, as described below.

**Fig. 1.**
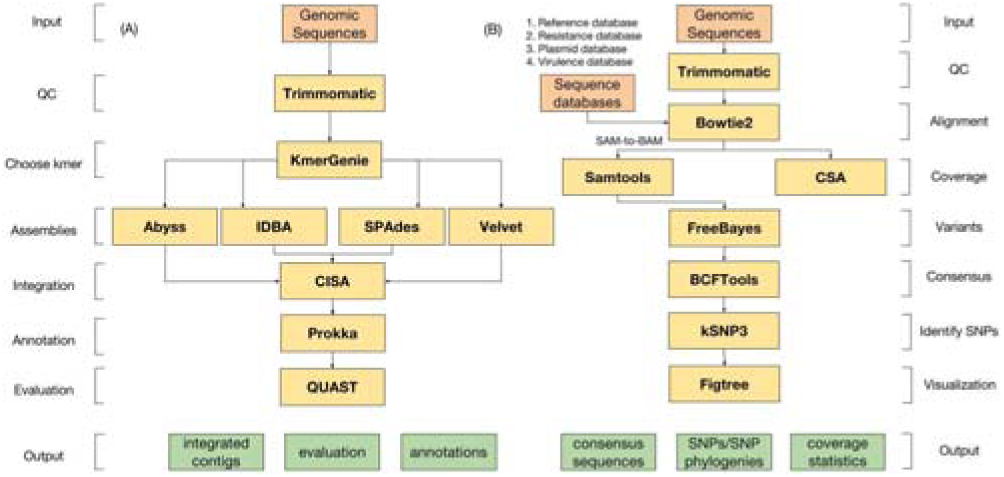
Implementation diagram for each step in the assembly and alignment modules (inputs, tools utilized, and outputs).

### 2.1 Assembly Module

Because no single assembler consistently produces an optimal genome assembly (Bradnam et al., 2013) in isolation, we utilize the results of four *de novo* genome assemblers to construct a consensus assembly of higher contiguity and accuracy. This module also supports methods for identifying and classifying genomic features of interest, a process called annotation, as well as reporting on common quality score metrics for each resulting assembly.

Input to the Tychus assembly module consists of a collection of paired fastq files. These files are preprocessed using Trimmomatic (Bolger, Lohse, & Usadel, 2014) to remove adapter sequences, low quality base pairs and sequence fragments. This prevents adapter sequences from being used to build the resulting assemblies, and has been shown to improve the quality of *de novo* assemblies as measured by their N50 statistic (2014). Preprocessed reads are then assembled with Abyss (Simpson et al., 2009), IDBA-UD (Peng et al., 2012), SPAdes (Bankevich et al., 2012), and Velvet (Zerbino & Birney, 2008). Prior to *de novo* assembly, an important step is to choose a *k*-mer value with which to build the underlying De Bruijn graph. As Abyss and Velvet do not iterate on multiple *k*-mer values, KmerGenie (Chikhi & Medvedev, 2014) is used to optimize this value prior to assembly. Contigs produced from each assembler are combined and integrated using CISA (Lin & Liao, 2013) to construct a super (consensus) assembly of higher quality and contiguity. Prokka (Seemann, 2014) is then used to annotate the super assembly. Lastly, contigs are evaluated with QUAST (Gurevich et al., 2013) to evaluate various common assembly score metrics, such as number of contigs, largest contig, and N50.

### 2.2 Alignment Module

The Tychus alignment module takes in six inputs: a collection of paired-end fastq files, a fasta-formatted reference genome, and optional databases of plasmid, resistance, virulence factor, and draft genome nucleotide sequences. Similar to the assembly module, reads are first quality-filtered using Trimmomatic (Bolger, Lohse, & Usadel, 2014), which helps to decrease the number of misalignments downstream. Next, Bowtie2 (Langmead & Salzberg, 2012) is used to align reads to the user-input reference genome, as well as the plasmid, resistance, and virulence databases. The results from alignment to the resistance, virulence and plasmid databases are used by an in-house tool to determine the overall coverage of each feature (plasmids, virulence factors, and antimicrobial resistance genes) present in each sample, with higher coverage features reducing the number of false-positive gene identifications. Files resulting from alignment to the reference genome are utilized by Freebayes (Garrison & Marth, 2012) to identify sequence variants and single nucleotide polymorphisms (SNPs), which are utilized by BCFtools (Li, 2011) to obtain consensus sequences for each sample. These consensus sequences, in addition to the optional genomes produced from the assembly module, act as draft genomes, which can be used by kSNP3 (Gardner, Slezak, & Hall, 2015) to identify related SNPs and build SNP phylogenies. The Newick formatted phylogenies produced by kSNP3 (2015) are then rendered into a user-defined image format using the command-line version of Figtree (https://github.com/rambaut/figtree).

## 3 Case Study

We used 15 *L. monocytogenes* samples (acc. no. PRJNA374745) with a reference genome (acc. no. PRJNA61583) and a 64-core (AMD Opteron Processor 6378) Ubuntu server to provide results for a common use case for each Tychus module. By default, all 15 samples were run in parallel using all available computing cores.

The assembly module was run with default parameters for each sample. Assemblies, annotations, and all assembly summary statistics were computed in under an hour, using approximately 40 gigabytes (GB) of RAM. Consistent with previous findings (Lin & Liao, 2013) the CISA integrated contigs produced superior assemblies in terms of N50, contig length, and number of contigs in nearly all samples (see Table 1).

**Table 1.**
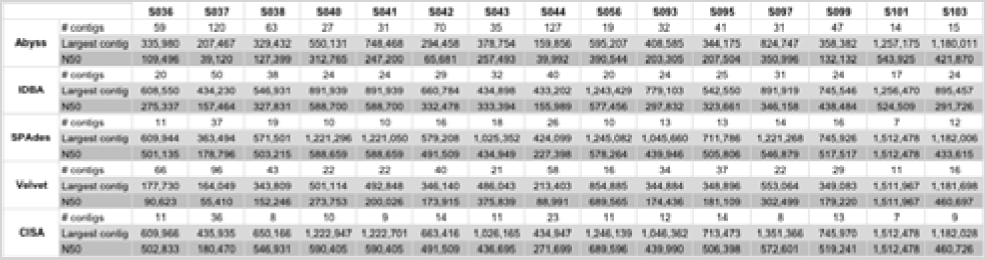
Assembly results for 15 *L. monocytogenes* samples.

Similarly, the alignment module was run with default settings for each sample. All sequence alignments, consensus sequences, SNPs, SNP phylogenies, and phylogenetic trees were produced in under 30 minutes using approximately 18 GB of RAM. A total of 77,601 SNPs were identified from all 15 strains, with 36,503 of these identified as core (SNPs present in all genomes); 41,098 identified as non-core; and 56,894 identified as majority SNPs present in a user-defined fraction of all samples (default 0.75).

In the end, it took Tychus <2 hours and <41GB of RAM to turn raw WGS data for 15 samples into high-quality assemblies with complete annotation and robust phylogenetic trees.

## Conclusion

Tychus is built upon existing open-source virtualization technology and provides a pipeline framework that is intuitive and easy-to-use for both novices and developers. It takes advantage of emerging multi-core computing architectures and large memory servers to deliver results quickly and reliably without the need for extensive bioinformatics or computing expertise. Though Tychus can be used on modest machines (with a penalty to run time), it is intended for use on servers with large amounts of RAM and disk space and with multiple computing cores.

## Acknowledgements

*Funding:* This work has been supported by startup funding from Colorado State University provided to ZA.

*Conflict of Interest:* none declared.

